# Neural dynamics of updating social impressions during movie watching

**DOI:** 10.64898/2026.05.13.724907

**Authors:** Jin Ke, Rhea Madhogarhia, Marvin M. Chun, Monica D. Rosenberg, Yuan Chang Leong, Hayoung Song

## Abstract

Updating impressions of others is essential to navigating social life. As we get to know an individual, we update our impressions of them in accordance with new information. How the brain dynamically revises impressions in real time under naturalistic conditions remains unclear. Here, we address this question using functional magnetic resonance imaging (fMRI) and natural language analysis in a naturalistic social cognition paradigm. Across 10 runs, participants viewed a character-driven TV episode, reported moments of insight, and described their impressions of the characters. Results reveal that impressions progressively evolved over time. Individuals with more similar existing impressions exhibit greater neural synchrony during movie-watching, which predicts convergence in post-movie impressions. Neural synchrony in the right superior temporal sulcus (STS) mediates the influence of initial similarity on later alignment. Insight moments accompany neural pattern shifts widespread across cortical regions, including the temporoparietal junction (TPJ), dorsomedial prefrontal cortex (dmPFC) and STS, and the magnitude of the shifts tracks the degree of impression updating. Specifically, distinct forms of insight selectively update complementary components of impressions: character insight updates person-centered representations, whereas non-character insight shapes social-event structure. Together, these findings show that people update their impressions of others through dynamic shifts in distributed brain activity patterns at moments of insight, providing a novel and ecologically grounded neural account of social cognition.

**Significance statement:** People do not simply form first impressions and keep them. As we learn new things about others, our judgments change—sometimes gradually, sometimes in a flash of insight. By tracking brain activity while people watched a TV show and described how their impressions of the characters evolved, we found that people who saw a character in more similar ways also show more similar brain responses when watching them onscreen, which in turn leads to convergence in subsequent impressions. Such updating is closely tied to “aha” moments, when a person is suddenly understood in a new way. These moments were marked by rapid shifts in the brain representational patterns, and larger shifts predicted greater changes in impressions. These findings offer a more naturalistic account of how the brain revises our understanding of other people in everyday life.

## Introduction

We navigate the social world by forming and continuously updating impressions of other people. Through everyday interactions, we rapidly infer what others are like. These impressions guide how we interpret their minds and choose to act toward them^1–3^, shaping affiliation and avoidance, trust and distrust, and decisions with lasting consequences—from romantic attraction and election to stereotypes and punishment^4–14^. Yet, impressions are not static. As new social information becomes available, earlier judgments can be revised when a new realization reshapes how the others are perceived.

Prior studies have identified the cognitive processes and neural systems involved in forming and updating social impressions. Studies of first impressions demonstrated that perceptual cues in faces^15–18^, voices^19–21^, and bodies^22–24^ can bias initial trait judgements and have identified how the brain supports forming impressions based on others’ behavior^25–30^. This work linked impression formation consistently with the dorsomedial prefrontal cortex (dmPFC), posterior cingulate cortex (PCC) and amygdala. Other studies have examined how these impressions change when new information confirms or conflicts with prior evaluations^25,31–37^, and have shown increased activity in the superior temporal sulcus (STS), temporal parietal junction (TPJ), rostrolateral PFC and PCC in this process. While this work has established where social impressions are expressed in the brain, it remains less clear how these neural representations change over time, and how they reflect individuals’ evolving impressions.

One possibility is that impression updating unfolds through gradual accumulations of evidence, with internal models of others revised continuously throughout the course of social perception. In this process, rather than replacing one judgement with the next, perceivers carry forward prior impressions as provisional models of others—mental priors that shape how subsequent behavior is interpreted. New information is thus integrated not in isolation, but in relation to what is already inferred about the person^33,38,39^. These evolving judgments emerge from an interplay between bottom-up social cues and top-down influences carried by the perceiver, including affect, attitudes, and personality^16,40^. For instance, an initial impression of a colleague as warm and trustworthy can bias how later behavior is processed: when they interrupt someone, this cue may be downweighted or reinterpreted during online social perception, yielding an updated impression that preserves the prior. Yet, a fine-grained characterization of how impressions are implemented in the brain over time—how they shape neural responses to new social information, are dynamically integrated, and give rise to updated impressions—remains lacking.

An alternative, complementary possibility is that impression updating is driven not only by gradual accumulation, but by discrete moments in which understanding shifts abruptly. For example, learning a hidden motive behind someone’s behavior can abruptly reframe prior interactions, leading to a sudden revision of their character. Such moments, often described as insight, are marked by sudden change in the representations of a stimulus, event or problem^41–43^. Prior work has studied insight predominantly in non-social domains, demonstrating that these changes in understanding are accompanied by reconfiguration in neural representation patterns^43–45^, for instance, during problem solving^44,45^, memory retrieval^46–48^ and narrative comprehension^48,49^. Extending this view to social cognition, impression updating may happen through moments of insight: new information about another person reshapes internal models of others, reflected as a sudden shift in neural representation patterns.

The central aim of the current study is to characterize the neural dynamics underlying impression updating and to determine the role of insight moments in this process. Addressing this question requires a setting where social cues unfold dynamically across time and contexts. Narrative stimuli, such as movies, are well-suited for this goal as they capture the complexity and richness of real-world social cognition through coherent, evolving storylines embedded in rich contextual information, while retaining experimental control^50–53^. In contrast, conventional paradigms often rely on static, decontextualized stimuli—such as isolated faces, trait words, or brief vignettes—thereby constraining ecological validity. In addition, prior research has largely depended on subjective ratings of predetermined trait dimensions, such as trustworthiness and dominance, yielding a low-dimensional characterization of social impressions. High-dimensional approaches, which represent impressions across many features (e.g., trait ratings or embeddings from natural language), may provide a more complete account of social impressions^54^. Here, we utilized natural language processing, neuroimaging, and computational analyses to assess the unconstrained, rich semantic meanings of social impressions through in-scanner speech.

In the current study, we collected fMRI data as participants (*N* = 36) viewed an audiovisual social narrative presented in a temporally scrambled order^48^. Over the course of 10 runs, participants watched segments of the movie while pressing an “aha” button to indicate moments of insight and explained them, and verbally reported impressions of each of the four main characters. We transcribed these verbal reports and used a text embedding model to capture the semantic meanings of social impressions. We then performed an intersubject representational similarity analysis^55^ to test whether individuals with more similar prior impressions of a character showed more similar neural responses while later viewing them onscreen, and whether greater neural similarity in turn predicted convergence in subsequent impressions. We next tested whether neural synchrony during movie watching mediated the change in character impressions. Finally, by analyzing neural pattern shifts around “aha” moments, we tested whether their magnitude tracks the degree of impression updating and whether distinct insight types selectively update different components of social impressions—character insights updating person-centered representations and non-character insights updating social-event structure. Taken together, our findings reveal that social impression updating occurs through both the ongoing neural activity during narrative processing and transient neural pattern shifts at moments of insight.

## Results

### Social impression updating during narrative comprehension

Thirty-six individuals watched a 41-minute, 40-second segment from an audio-visual movie, the first episode of NBC’s *This Is Us*, while undergoing functional MRI^48^. Participants were screened to ensure that they had never watched any episode of the show before. We divided the movie into 48 scenes (each 52 ± 21 sec), scrambled their temporal order, and grouped them into 10 runs. By disrupting temporal coherence and assigning participants to one of three different scrambling orders, participants encounter character information at different times, increasing variability in when and how impressions are updated across individuals. In each run, participants watched the movie and pressed an “aha” button whenever they experienced a subjective moment of insight (i.e., as they understood something new about the show’s characters or events; Part I). We then presented movie screenshots at those exact moments of button presses and asked participants to verbally explain why they pressed the “aha” button (Part Ⅱ). Finally, participants verbally described impressions on each of the four main characters (Part Ⅲ; **Fig. 1a**). Specifically, we asked participants to talk about their thoughts about the character, and what they had learned about the character’s behavior, thoughts, feelings, intentions, and relationships with other characters.

**Figure 1.**
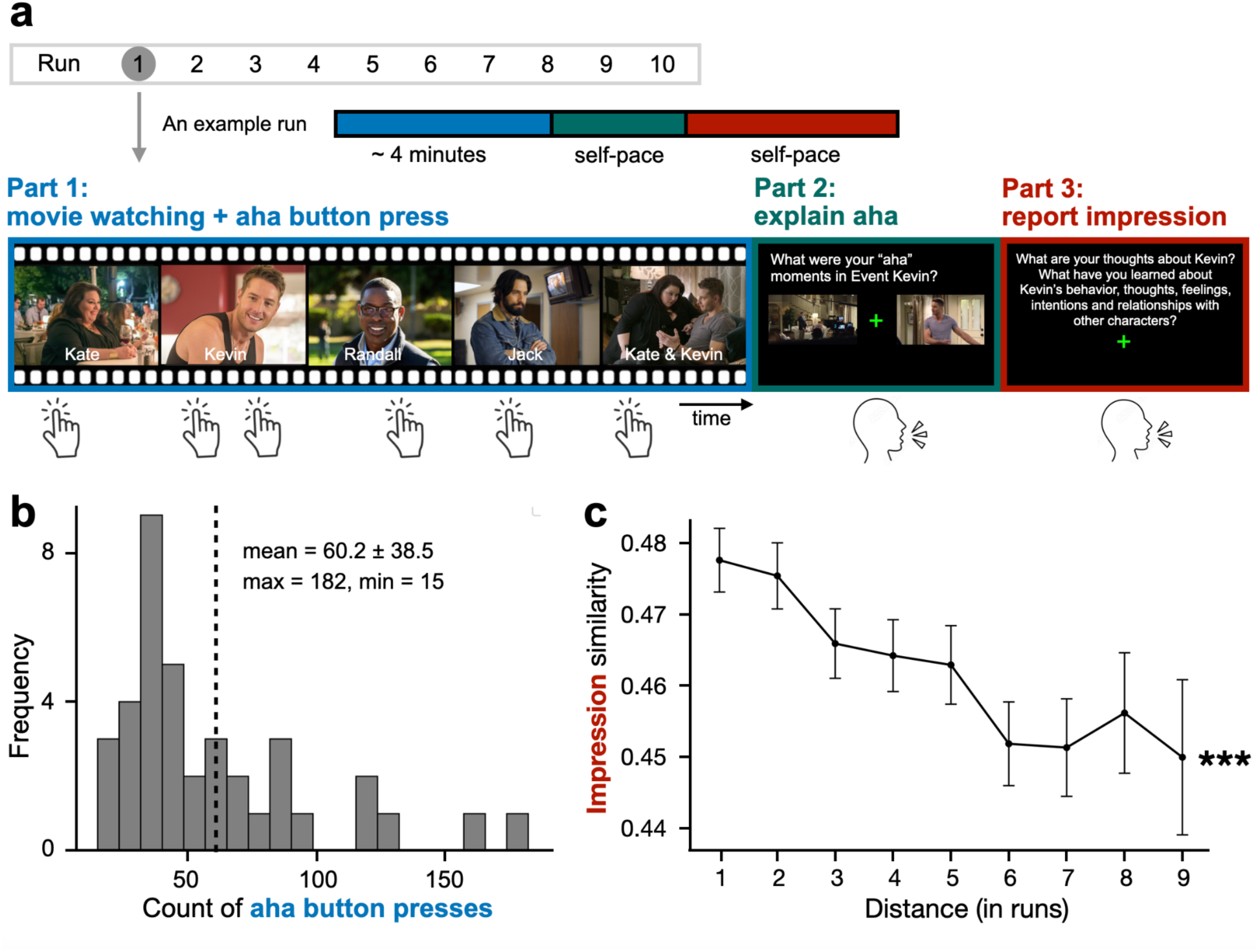
**(a)** Overview of the experimental design. 36 participants completed a ten-run fMRI session. In each run, they first watched temporally scrambled movie segments and pressed an “aha” button whenever they experienced a subjective moment of insight. Next, they viewed screenshots corresponding to those button presses and verbally explained their insights. Finally, they verbally described their impressions for each of the four main characters. **(b)** Distribution of “aha” button presses across participants. The x-axis shows the number of presses, and the y-axis indicates the number of participants. **(c)** Impression similarity decreases with increasing temporal distance. Impression similarity, plotted on the y-axis, was computed as the cosine similarity between 512-dimensional Universal Sentence Encoder sentence embeddings of participants’ reported impressions. Each black dot represents the mean similarity at a given temporal distance (x-axis, ranging from 1–9), and error bars represent 95% confidence intervals. ***: p < .001

We first examined participants’ reports of moments of insight during movie watching, as indexed by “aha” button presses. The number of “aha” presses ranged from 15 to 182 per individual (mean = 60.2, SD = 38.5; **Fig.1b**), indicating that participants frequently had new understandings while comprehending the social narrative, with substantial variability across individuals. Three independent annotators categorized participants’ open-ended explanations of their aha moments into eight topics (e.g., characters, current scene understanding, memory retrieval). Insight about the characters was one of the most frequent topics (mean = 45.8%, SD = 19.2%; **Suppl. Table. 1**), consistent with the character-driven nature of the stimulus.

Next, to investigate how social impressions change over time, we analyzed participants’ verbal descriptions of each character. Four participants were excluded due to missing verbal transcripts. The semantic content of these impressions was quantified as 512-dimensional sentence embeddings using Google’s Universal Sentence Encoder (USE), and the similarity between these embeddings was assessed using cosine similarity. The approach has been previously used to measure semantic similarity of narrative content^56–58^. We built nested mixed-effects models to predict between-run impression similarity based on their temporal distance (in runs), with intercepts of character, group, and subject nested within groups included as random effects. Results revealed that, within individuals, impressions became significantly less similar as their temporal distance increased (*β* = -.065, *s.e.* = .012, *z* = -5.214, p < .001; **Fig. 1c**), suggesting that participants’ thoughts about characters gradually changed throughout runs, with impressions of the nearby runs more similar than distant runs.

Together, these findings suggest that the majority of the reported insights were about characters and participants gradually revised impressions of the characters as the narratives unfolded. In subsequent analyses, we asked how the brain encodes these evolving impressions and whether neural dynamics accompanying insights signal impression updating.

### Shared neural responses track shared impression of others

To test how the brain encodes social impressions, we performed an intersubject representational similarity analysis (IS-RSA) within each scrambling-order group, relating impression similarity with neural synchrony during movie watching (**Fig. 2a**). Seven participants were excluded due to excessive head motion (*N* = 3; >10% of frames censored during movie viewing) or missing verbal transcripts (*N* = 4). We assessed pre-movie impressions—verbal reports from the preceding run—to test whether similar prior impressions of others predict similar neural activity when later watching them onscreen. We also assessed post-movie impressions—verbal reports in the current run—to examine whether similar neural activity during movies predicts similar subsequent impressions.

**Figure 2.**
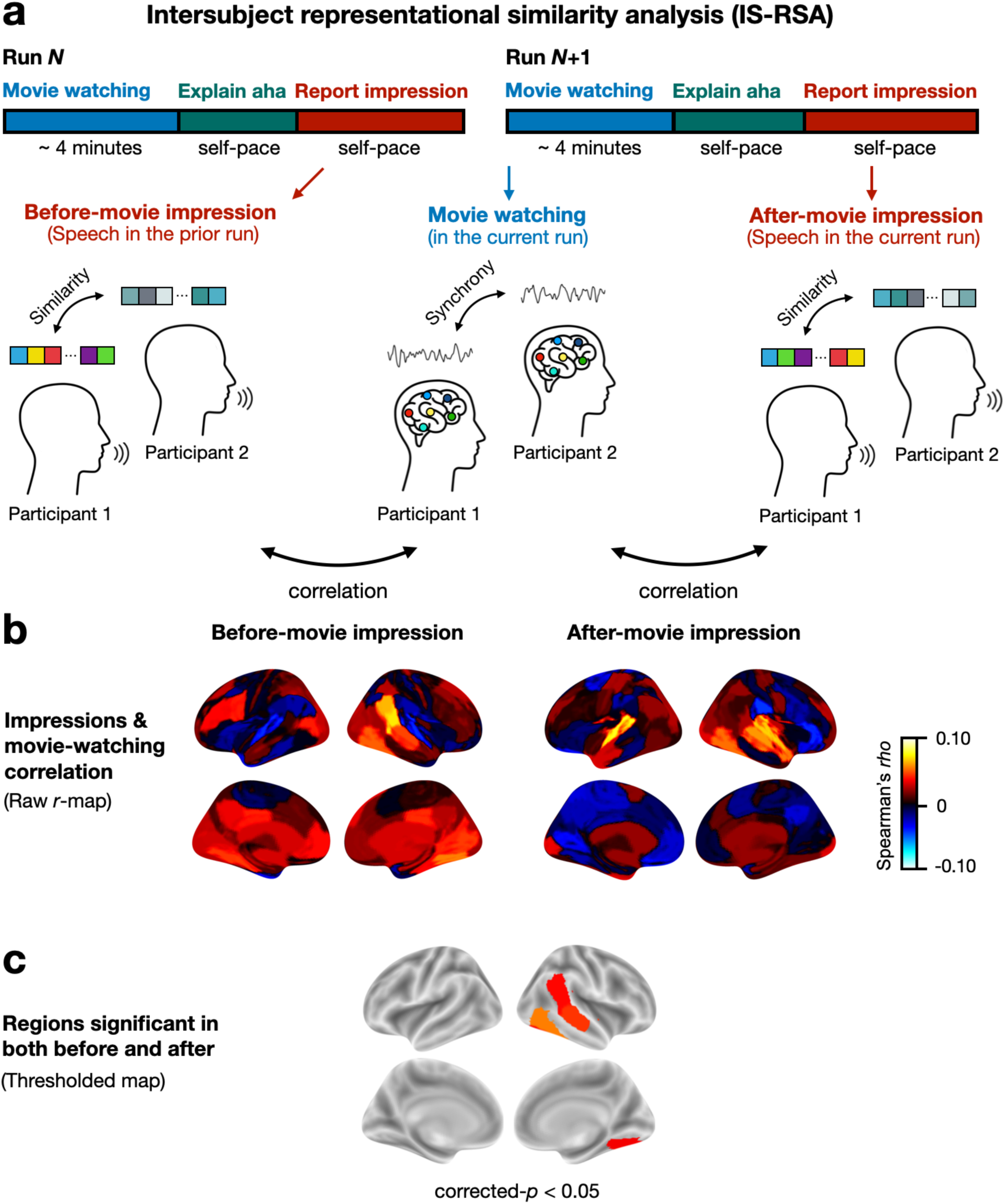
Linking shared impressions to synchronized neural responses. **(a)** Intersubject representational similarity analysis (IS-RSA). For each ROI, neural synchrony during movie-watching was computed as the Pearson correlation between BOLD activity time series from every pair of participants within groups. Before-movie impressions were defined as reported impressions in the preceding run, and after-movie impressions as those from the current run. Between-individual impression similarity was quantified as cosine similarity between their USE embeddings. We related neural synchrony with impression similarity using Spearman correlation. **(b)** Brain maps show the Spearman correlations between neural synchrony and impression similarity before (left) and after (right) movie watching. **(c)** Regions whose neural synchrony tracks both before and after-movie impressions. The log-transformed *p*-statistics from the “before” and “after” IS-RSA analysis were combined into a single chi-squared statistic following Fisher’s method. Only ROIs showing significant effects in both were plotted (FDR corrected-*p* < 0.05). The colors indicate the Spearman’s *r*-values averaged across the two conditions (i.e., before and after).

Impression similarity was quantified as the cosine similarity between the USE embeddings of transcribed verbal descriptions. While prior work has implicated specific regions in impression updating, social representations may be distributed across the brain. To capture this distributed structure in an unbiased manner, we parcellated the fMRI data using a functional brain atlas consisting of 116 regions of interest (ROI) covering 100 cortical^59^ and 16 subcortical regions^60^. For each brain region, neural synchrony was computed as the Pearson correlation between the voxel-averaged time series of two individuals who watched the movie in the same sequence. We correlated the matrices of between-participant impression similarity and neural synchrony for each scene and compared the resulting distribution of Fisher-*z* transformed *r*-values against zero with bootstrapping (10,000 iterations). We assumed a two-tailed significance test and corrected for multiple comparisons.

We found that dyads with more similar impressions of the characters before watching the movie events exhibited greater neural synchrony when watching them onscreen in regions including the right temporoparietal junction (TPJ), angular gyrus, superior temporal sulcus (STS) and inferior temporal (IT) cortex (FDR-corrected *p* < 0.05; **Fig. 2b**, left). Higher neural synchrony during movie watching, in the bilateral medial temporal gyrus (MTG), right STS, superior temporal gyrus (STG), and the right auditory cortex and IT cortex, predicted more similar impressions after movie-watching (FDR-corrected *p* < 0.05; **Fig. 2b**, right; See **Suppl. Fig. 1** for the thresholded brain map respectively for before and after). To identify regions in which neural synchrony tracked both before- and after-movie impressions, we combined evidence across the two IS-RSA analyses using Fisher’s method, which sums the log-transformed *p*-values from regions with consistent effect directions into a single statistic evaluated against a chi-squared distribution (see Methods). Neural synchrony in the right TPJ, STS, and IT cortex tracked both prior and subsequent impression similarity (FDR-corrected *p* < 0.05; **Fig. 2c**), suggesting that similar social impressions are positively associated with similar neural responses during movie watching. Together, these findings suggest that social impressions are encoded across social-cognitive and sensory regions and validate that USE-embeddings capture the semantic meanings of social impressions.

### Neural synchrony in the right superior temporal sulcus mediates impression updating

We further examined whether neural synchrony during movie watching shapes the relationship between pre- and post-movie impression similarity (*N* = 29). We built causal mediation models, which estimate how much of the predictor’s causal effect on the outcome is transmitted through the mediator (**Fig. 3a**). Separate models were fit for each ROI, with multiple-comparison correction applied across 116 ROIs.

**Figure 3.**
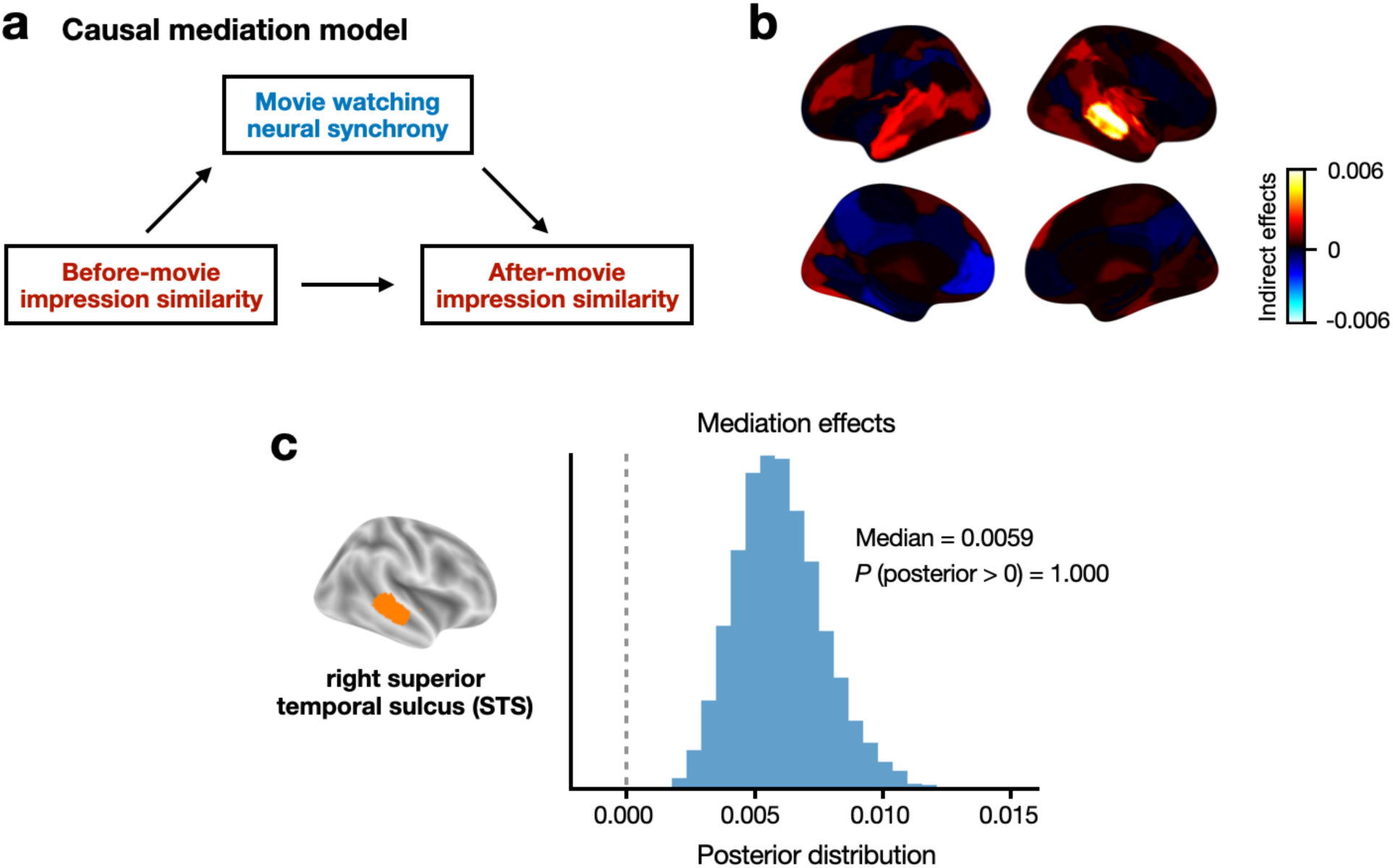
Shared neural responses in the right superior temporal sulcus (STS) mediate the relationship between pre- and post-movie social impressions. **(a)** Schematics of the causal mediation model. **(b)** Brain maps showing the indirect effect of neural synchrony on the relationship between pre- and post-movie impressions. **(c)** The posterior distribution of the mediation effects in the right STS.

The total effect of before-movie impression similarity on after-movie similarity was significant (total effect = 0.0783, *p* < 0.001), suggesting that pre-existing impressions serve as mental priors that shape subsequent impressions. Mediation analysis identified an indirect pathway through neural synchrony in the right superior temporal sulcus (right STS; ACME = 0.006, corrected-*p* < 0.001; **Fig. 3b**, and see **Fig. 3c** for the posterior distribution of the mediation effect), accounting for 7.7% of the total effect. The direct effect remained significant (ADE = 0.072, p < 0.001), indicating partial mediation, whereby shared neural responses in the right STS contribute to—but do not fully account for—the influence of initial similarity on later convergence of impressions. Before correction, additional regions—including the left precuneus, MTG, visual cortex, and the right TPJ, MTG, and dlPFC—showed significant or marginally significant mediation effects (**Suppl. Fig. 2**; **Suppl. Table 3**), although none survived corrections for multiple comparisons. These findings suggest that individuals with more similar existing impressions exhibited greater neural synchrony in the right STS during movie-watching, which in turn predicted convergence in post-movie impressions. This thus identifies right STS as a central brain region through which shared initial impressions are transformed into shared subsequent impressions.

### Neural pattern shifts occur at moments of insight and reflect impression updating

Having established that neural responses during movie watching track the evolving impressions across individuals, we proceeded to investigate whether certain moments are especially crucial for these changes (*N* = 29). Prior work on insight suggests that abrupt shifts in neural activity patterns reflect changes in mental models and updating of ongoing interpretations^45–49^. Thus, we hypothesized that moments of character insight—as participants arrive at a sudden new understanding of a character—would be reflected as transient shifts in neural activity patterns, and that larger shifts would be associated with greater changes in impressions. To isolate the contribution of insight moments above and beyond regular ongoing processing during movie watching, in the following analysis we used non-insight moments as the baseline, defined as time points outside the −5 to +3 TR window surrounding any aha button press.

To test whether insight is accompanied by rapid updating of neural representations, we quantified moment-to-moment neural pattern shift as 1 minus Pearson correlations of voxel patterns in the rSTS between two consecutive TRs. These shifts were computed for each TR and aligned to aha events, yielding a time-resolved measure from −5 to +3 TRs relative to each button press. The hemodynamic lag was accounted for by temporally shifting the neural response time courses by 4 TRs (TR = 1.2s). If a participant reported multiple aha moments for the same character within a movie segment, we averaged the corresponding neural pattern shifts to obtain a single estimate for that character in that run.

We assessed neural pattern shifts near character-related, non-character, and general insight, using non-insight moments during the same movie-watching run serving as the baseline condition. Character-related insights—defined as moments reflecting understanding of a character’s traits, behavior, thoughts, or feelings (**Suppl. Table 1**)—accounted for 45.8% of all aha events (≥2 of 3 raters). Non-character insights, identified when none of the three raters attributed the moment to character-related insights, comprised 33.9%. The remaining events, with only a single rater endorsing a character interpretation, were excluded from analysis due to insufficient inter-rater agreement, and thus percentages do not sum to 100%. General insight referred to all moments in which participants pressed the aha button, regardless of category. In the right STS, the region we identified to link prior and subsequent impressions in the earlier mediation analysis, neural pattern shifts during character insight were significantly above the non-insight baseline from 2TRs before to 1TR after the aha button press, whereas non-character insight and general insight exhibited no such effects (**Fig. 4a**). This indicates that neural representations in the right STS are more likely to shift around moments of character insight than other insight types, consistent with its role in social impression processing. This provides initial evidence that neural pattern shifts near reported insight moments may reflect updates in character representations, motivating a direct test of whether the magnitude of these shifts tracks the degree of impression change.

**Figure 4.**
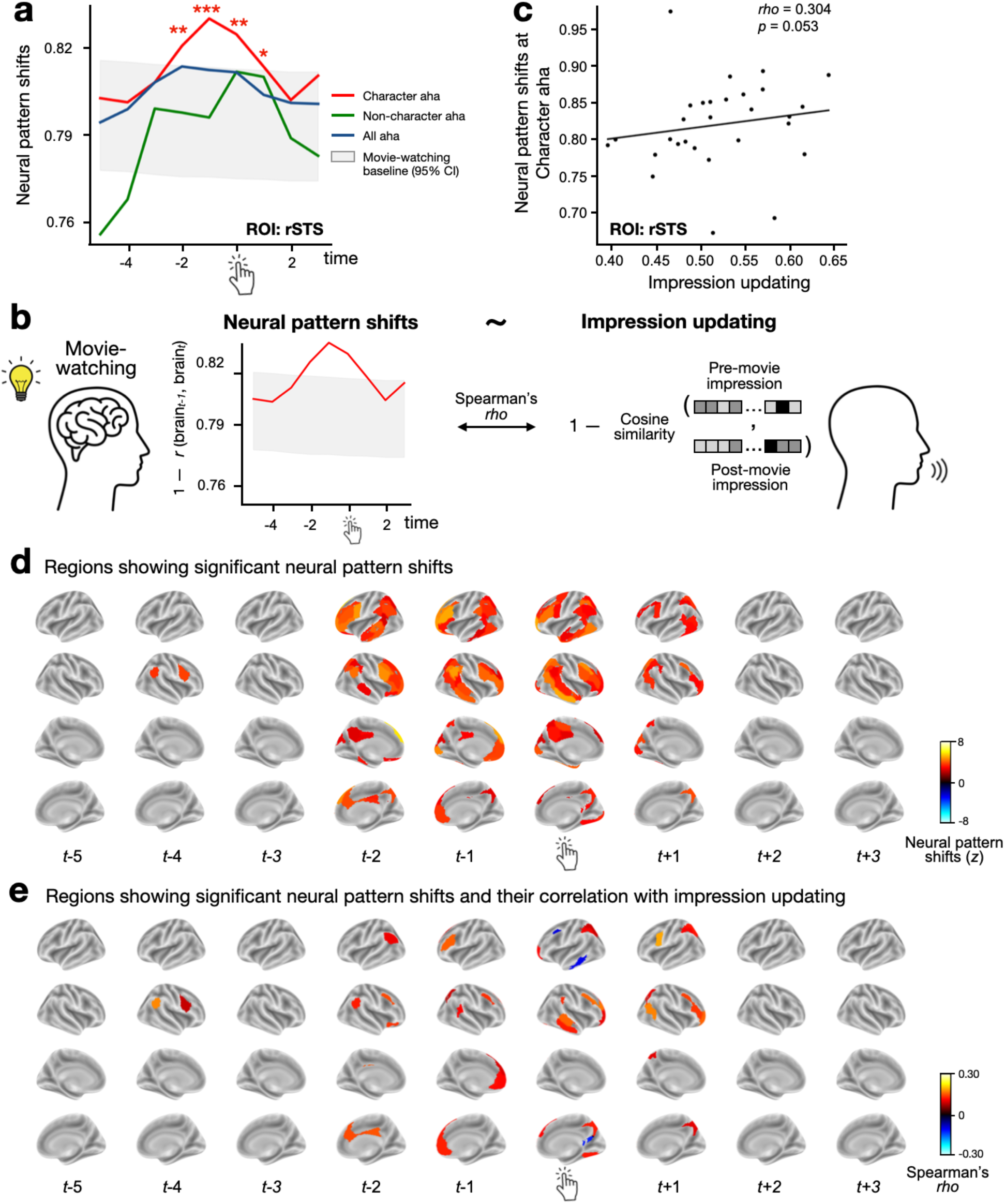
Neural pattern shifts occur near moments of character insight and track impression updating. **(a)** Neural pattern shifts during moments of character insight, non-character insight, and general insight in the right STS (rSTS). The gray area represents a null distribution of neural pattern shifts from non-insight moments. **(b)** Schematics of correlating neural pattern shifts, measured as one minus the Pearson correlation between voxel patterns in consecutive TRs, with impression updating, measured as one minus the cosine similarity between USE-embeddings of pre- and post-movie impressions. **(c)** Correlation between neural pattern shifts at character aha in the rSTS averaged from -2 to +1 TR and impression updating. Each dot represents cross-participant correlation for each scene. **(d)** Whole-brain neural pattern shifts as a function of time relative to character-related aha button presses. Brain maps show z-statistics that compare neural activity patterns near aha moments against a null distribution derived from non-aha moments. Significance was corrected for multiple comparisons across 116 ROIs and 9 TRs. **(e)** Regions showing both significant neural pattern shifts and their correlation with impression updating. Significance was corrected for multiple comparisons across 116 ROIs and 9 TRs. *: *p* < 0.05, **: *p* < 0.01, ***: *p* < 0.001.

To this end, we correlated the magnitude of these neural pattern shifts to changes in impressions, quantified as 1 minus cosine similarity between the USE-embeddings of pre- and post-movie impressions (**Fig. 4b**), across participants. Specifically for the right STS, the resulting correlations were significantly above zero across −5 to +3 TRs relative to aha button presses (paired-*t* = 6.492, *p* < 0.001). The observed effects remained significant when compared against the 95th percentile of the more conservative null distribution derived from non-insight moments in the same movie-watching run (paired-*t* = 2.697, *p* = 0.014; **Suppl. Fig. 3**). The correlations did not peak at specific moments near aha button presses, but were generally high throughout -5 to +3 TRs compared to non-insight moments. To summarize the overall contribution of the right STS to impression updating, we averaged neural pattern shifts within the window showing reliable reconfiguration (−2 to +1 TR; **Fig. 4a**) and related this to impression updating. This analysis yielded a marginal effect (Spearman’s *rho* = 0.304, *p* = 0.053; **Fig. 4c**), indicating that neural pattern shifts in the right STS may relate to impression updating preferentially at moments of character-related insight rather than more generally during movie-watching.

The marginal effect observed in the right STS motivated a whole-brain analysis to identify where neural pattern shifts reliably track impression updating across the brain. To this end, we repeated the analytical pipeline on each of the 116 ROIs, again quantifying neural pattern shifts as 1 minus TR-to-TR voxel pattern similarity. Replicating our previous findings^48^ using hidden Markov Models to infer boundaries of shifts in neural activity, we observed significant neural pattern shifts near character insight moments across distributed regions of the brain (**Fig. 4d**; *z*-statistics reflect deviation from the null distribution of non-insight moments; FDR-corrected for multiple comparisons across the 116 ROIs and 9 TRs). These effects peaked from 2 TRs before up to its occurrence, with 38 (-2 TR), 36 (-1 TR), and 43 (0 TR) cortical regions showing significant neural pattern shifts (**Fig. 4e**). No significant neural pattern shift was observed in subcortical regions. This finding suggests that transient neural pattern shifts in distributed areas of the cortex were observed near reported moments of insight about characters.

We further related the magnitude of neural pattern shifts with the degree of impression updating. For interpretability, we visualized ROIs that showed both reliable neural pattern shifts and significant associations between neural pattern shift magnitude and impression updating (corrected-*p* < 0.05; **Fig. 4f**; see **Suppl. Fig. 4 & 5** for results from non-character aha moments). We found that, near moments of character insight, the degree of neural pattern shifts in regions including TPJ, dlPFC, dorsomedial prefrontal cortex (dmPFC), ventromedial prefrontal cortex (vmPFC), precuneus, STS, and sensory regions positively correlated with the degree of impression updating. Notably, although neural pattern shifts were temporally selective around aha button presses—particularly from −2 to +1 TRs—their relationship with impression updating was not confined to this narrow window (**Suppl. Fig. 6**). This suggests that impression updating unfolds broadly across scenes surrounding insight, while still involving abrupt representational changes near the reported aha moments. Together, these findings suggest that social impressions are updated through discrete moments of insight, where transient shifts in neural representations occur.

### Character and non-character aha moments differentially shape impression updating

We reasoned that insights about characters should revise person-centered impressions—what a character is like, thinks, feels, or intends—whereas insights not centered on characters may instead revise understanding of the broader social event, such as what happened, why it happened, or how events and characters are related. We therefore asked whether these two insight types carried distinct semantic signatures and updated different components of social understanding.

First, as a validation of our operational definition of the two insight types, we trained a logistic regression model to classify character vs non-character from their USE-embeddings with 5-fold cross-validation and found reliable separability (AUC = 0.950 ± 0.010; **Suppl. Fig. 7a**; **Suppl. Fig. 7b** visualized the USE-embeddings in the Uniform Manifold Approximation and Project (UMAP)^61^ space), indicating differences in their semantic content.

We hypothesized that character and non-character insight diverge across two complementary representational measures. We defined the *person model* as a representation of an individual’s characteristics, including dispositional traits and mental states^62–64^. Language describing this model often shows an individual focus (e.g., pronoun use including she, her, he, him)^65,66^, reflecting attention to a specific person’s characteristics rather than external events. In contrast, we defined the *social-event* model to represent the broader situational context, including social relationships, group structure, actions and events, and the causal links between them^67^. Together, these two models distinguish between inferences about what a person is like and representations of what happens around them.

To examine how character and non-character insight moments differ, we analyzed the language participants used to describe these insights (*N* = 33). For each transcription, we computed the proportions of words assigned to each of the semantic category dictionaries (**Suppl. Table. 4**), constructed based on prior work on trait-related^64^ and mental state terms^68^. As expected, descriptions of character aha moments used significantly more words reflecting dispositional traits, mental states, and individual focus and fewer words related to social relationships, group focus, and event or action (Wilcoxon rank-sum test; all corrected-*p* < 0.001; **Fig. 5a**). We then averaged proportions of these categories within each model to derive summary measures of the person model and the social-event model. Using this summary measure, we found that character insights exhibited greater weight on the person model (70.2%) than the social-event model (29.8%; Wilcoxon signed-rank *z* = 21.36, *p* < 0.001; **Fig. 5b**). Conversely, non-character insights had more percentage of social-event model (62.2%) than person model (37.8%; *z* = 9.48, *p* < 0.001). These results validate our model definition, such that character insight, as compared to non-character insight, preferentially engages the person model than the social-event model.

**Figure 5.**
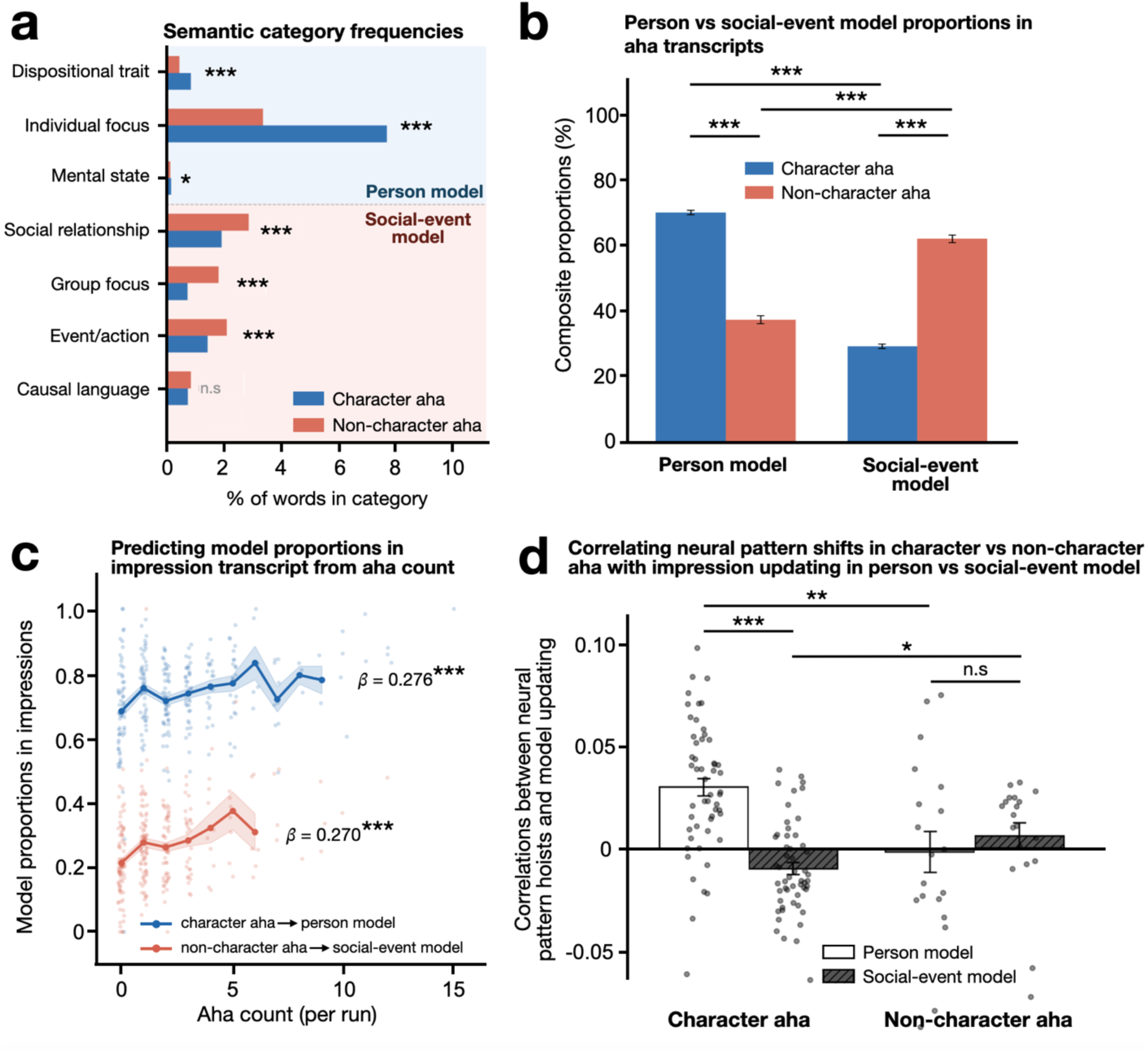
Character and non-character aha moments differentially shape impression updating. **(a)** Comparison between character and non-character aha on nine categories within the person and social-event model. The x-axis represents the percentage of words that belong to the category dictionary. **(b)** Comparison between the proportions of the person vs social-event models in character and non-character insights. The bars represent the group mean composite proportions, and the error bars represent standard error. **(c)** Nested mixed effect model predicting changes in impressions (person vs. social-event model proportions) from counts of aha button presses (character vs. non-character aha buttons). Each dot represents one run from one participant, the solid, larger dots represent the mean model proportions at each aha count, and the solid lines connect these dots. **(d)** Correlations between neural pattern shifts near character vs. non-character insights and impression updates in person vs. social-event models. Each dot represents a brain region that exhibited significant neural pattern shifts near character (51 regions) or non-character insights (19 regions). *: *p* < 0.05, **: *p* < 0.01; ***: *p* < 0.001, n.s.: not significant.

By analyzing the verbal reports of character impressions, we likewise observed a higher proportion of the person model (73.4% ± 21.0%), relative to the social-event model (26.6% ± 21.0%). This bias was consistent across ten runs and four characters (**Suppl. Fig. 8a-b**), despite clear clustering of their semantic content in UMAP space (**Suppl. Fig. 9**). Thus, a natural hypothesis is that the character-related insight moments would preferentially update the person model in social impressions, whereas non-character insights track updates in the social-event model.

We built nested linear mixed-effects models to predict impression updating from the number of aha button presses, with runs nested within subject and group included as a random effect. Impression updating was quantified as changes in the person vs social-event model proportions. We found that the more character insights participants experienced, the greater the updates in the person model (*β* = 0.270, *p* < 0.001), accompanied by corresponding decreases in the social-event model (*β* = -0.276, *p* < 0.001). In contrast, non-character insights exhibited the opposite pattern, being associated with greater updates in the social-event model (*β* = 0.276, p < 0.001) and corresponding decreases in the person model (*β* = -0.270, *p* < 0.001; **Fig. 5c**). These findings suggest that different forms of insight selectively reshape complementary components of social impressions.

Finally, we examined whether neural pattern shifts during character and non-character insights predict updates in the person and social-event model of impressions. In brain regions showing significantly higher neural pattern shifts near character insights (**Suppl. Fig. 4a-b**), these shifts preferentially tracked updates in the person model, showing stronger associations than with social-event model updates (*t* = 8.98, *p* < 0.001; **Fig. 5d**, left). By contrast, in regions showing significant pattern shifts near non-character insights (**Suppl. Fig. 4c-d**), correlations with the social-event model updates were numerically higher than with person model updates but did not reach significance (*t* = 0.698, *p* = 0.494; **Fig. 5d**, right). These results suggest that neural pattern shifts selectively track updates to person-centered impressions during character insights, but exhibit no reliable selectivity during non-character insights. Together, our findings suggest that social impressions are updated through moments of insight accompanied by transient shifts in neural representations.

## Discussion

How we understand other people in everyday life fluctuates over time^40^. As new social evidence unfolds, we interpret it in light of prior beliefs and update our internal models of others. Yet, how the human brain revises these impressions over extended timescales in naturalistic settings remains unclear. In this study, we combined fMRI, natural language analyses, and a naturalistic social cognition task to characterize the neural dynamics of impression updating. Individuals who shared more similar initial impressions showed greater neural synchrony when later viewing the same characters, which predicted greater convergence in subsequent impressions. Mediation analyses identified the right superior temporal sulcus as a key neural hub through which shared initial impressions were transformed into shared later impressions. Furthermore, moments of character insight—when individuals had sudden new realizations of the characters—were accompanied by transient neural pattern shifts across distributed regions in the cortex, and larger shifts predicted greater subsequent impression updating. Character insights were more associated with updates to person-centered representations, whereas non-character insights were more associated with updates to social-event structure. Together, these findings reveal the neural dynamics of real-world social impression updating and identify character insight as a core mechanism through which impressions change.

Our findings extend neuroscientific investigations of social impressions from isolated stimuli to complex, temporally unfolding narratives, which are central to how people interpret experiences, exchange knowledge and form social connections^53,69,70^. Prior work has typically used brief cues or behavioral descriptions that either confirmed or violated an initially formed evaluation, and tested how neural activity tracks the incorporation of congruent or incongruent information into impression^25,31,33–35^. This work has identified a network—including dmPFC, PCC, STS, TPJ and amygdala—engaged during impression formation and updating. However, because these paradigms rely on discrete, experimenter-defined updates, they characterize how impressions change in response to controlled inputs. This leaves open how updates arise spontaneously in naturalistic contexts, where evidence accumulates across time in a more nuanced manner and must be interpreted against prior beliefs. By embedding impression updating within a rich narrative stream, our study shows that social impression updating is not reducible to isolated, one-shot events but a continuous, context-dependent process of internal model revision during ongoing social experiences.

The current results contribute to a rich literature suggesting human perception and cognition are shaped not only by sensory input, but also by prior beliefs, motivations and expectations^71–78^. Parallel work also shows that narratives likewise reveal structured individual differences: people with different beliefs, traits, or interpretations often exhibit more idiosyncratic neural responses to the same narrative despite the identical sensory input^79–83^. Our study extends this framework to social cognition by showing that neural responses to other people are shaped by the perceiver’s prior impressions. Using IS-RSA, we found that individuals with more similar prior impressions also exhibited more similar neural responses during subsequent social perception, and ultimately converged on more similar later impressions. These findings suggest that existing impressions act as mental priors that shape how new social information is processed over time.

While traditional IS-RSA establishes a correspondence between shared impressions and shared neural responses, the mediation analysis extends this framework by statistically examining a directional relationship linking prior impressions, neural synchrony during social perception, and later impressions. Specifically, neural synchrony during the processing of new social information mediated the association between initial and subsequent impressions, consistent with a pathway in which prior impressions shape ongoing neural processing, which in turn relates to later social understanding. In this way, the mediation framework advances IS-RSA beyond mapping representational similarity by identifying a candidate neural pathway through which impression updating unfolds over time.

Our findings identified the right superior temporal sulcus (STS) as a region mediating impression updating. This brain region has been long implicated in diverse aspects of social perception and cognition, including perceiving biological motion, faces, voices, social interactions, as well as understanding others’ actions, emotion, and mental states^84–95^. In contrast, the left STS has been shown to be involved in speech- and language-related processing, including phonetic analysis and audiovisual speech integration^96–99^, suggesting a relative hemispheric asymmetry in which the right STS preferentially supports social perception^100^. Consistent with this account, our results further extend its functional role by suggesting that the right STS not only represents socially relevant signals but also links prior impressions to the ongoing interpretation of unfolding social information. In our data, between-individual neural synchrony in this region linked shared prior impressions to later convergence in judgment, pointing to the right STS as a candidate region through which prior beliefs shape subsequent social understandings in naturalistic settings, in line with the predictive accounts of social cognition^101,102^.

One key theoretical contribution of this work is to situate social impression updating within the framework of insight. Insight has been characterized as the sudden emergence of a coherent interpretation through reorganization of previously weakly connected, ambiguous or conflicting information^41–43^, often accompanied by abrupt shifts in neural representations^45–49^. Extending this account to social cognition, our findings indicate that updating impressions of others is not solely a gradual accumulation of evidence, but can occur through discrete, insight-like transitions in understanding. In naturalistic contexts, new information can retrospectively reshape prior interpretations, updating neural representations of earlier events and revising internal models^47^. Consistent with this account, moments of character insight were accompanied by widespread neural pattern shifts around the time of reported insight, while the magnitude of these shifts tracked impression updating mainly in regions supporting social inference, including the STS, TPJ, dmPFC, and dlPFC.

While prior work showed increased activation of these regions during impression updating^25,33,35^, our results instead highlight changes in multivariate activity patterns, revealing that these regions dynamically reconfigure their representations at moments of character insight. The magnitude of this pattern shift scales with the degree of impression updating, suggesting that insight involves restructuring representations to incorporate new information into a coherent interpretation of others. Moreover, this reconfiguration is not confined to one single brain region but unfolds across a distributed social-cognitive network, echoing a growing recognition that social understanding emerges from coordinated interactions between brain regions^103–106^. Together, our work suggests a network-level mechanism in which transient shifts in neural representational patterns enable rapid updating of social impressions, linking moments of insight to lasting changes in how others are understood.

We note three potential limitations of the current study.

First, because we used a single narrative film, some effects may reflect properties specific to that story. Although the film was presented in three scrambled temporal orders—arguing against a simple effect of plot sequence—all groups were built from the same underlying narrative. We therefore cannot fully dissociate the observed effects from features of this particular story. Future work should test whether these findings generalize across diverse narratives.

Second, although participants’ speech reports captured rich and unconstrained accounts of insight during narrative comprehension, our classification of insight types was subjective. In the absence of established frameworks, we derived these categories inductively from participants’ verbal responses. While these categories appeared meaningfully related to impression updating, future work will be needed to define narrative insight types more systematically and to clarify their distinct roles in social cognition.

Third, the multitasking demands of the paradigm may have perturbed the natural flow of social perception. Specifically, requiring participants to report insight via button presses during movie viewing may have altered ongoing processing. Although prior work suggests that minimal responses during naturalistic tasks do not substantially disrupt behavior or neural dynamics^107^, it remains unclear whether this conclusion extends to such online reports. Future studies should therefore test whether the present effects replicate without concurrent button presses, for example by examining whether similar relationships between neural pattern shifts and impression updating are observed in the absence of overt responses.

Despite these caveats, the current study reveals the neural dynamics that support impression updating in real-world settings and identifies moments of insight about others’ traits, intentions, or mental states as a key component of this process. Our findings suggest that impression updating reflects the impact of prior beliefs on the interpretation of unfolding social information, particularly during moments of insight that reshape understanding. Our work lays the foundation for future investigations into how the brain updates internal models of others during naturalistic social experience.

## Methods

### Experimental procedure

We analyzed openly available preprocessed fMRI data collected by our research group^48^. Participants (*N* = 36) watched a temporally scrambled version of a 41-minute, 40-second video from the first episode of NBC’s *This Is Us* while undergoing functional MRI. This particular television episode was chosen as it features a multi-threaded narrative involving four main characters whose storylines unfold independently and converge into a big reveal, providing rich social dynamics and thus suitable for examining impression updating. We segmented the episode into 48 scenes based on the director’s cut, where most scenes feature only one of the main characters, Kevin, Kate, Randall, or Jack (**Suppl. Table 2**). We then grouped these scenes into nine blocks in scrambled sequences (five scenes each block), with the last three scenes showing the big reveal forming a final block. These blocks were again scrambled into three different viewing orders (N=12 per scrambled-order group) such that participants in different groups watched the episode in different sequences. This allowed us to compare the effect of knowledge and contexts while keeping the perceptual input constant. Moreover, temporal scrambling of the narrative makes comprehension more challenging and induces more dynamic social impression updating.

The 36 participants were randomly assigned to one of the three groups. Before scanning, they were informed that the episode was segmented into multiple scenes and scrambled in their temporal order. They were informed that the sequence was intentionally shuffled to elicit dynamic fluctuations in comprehension, so that they would experience multiple “aha!” moments as they connect the dots to understand the narrative plot. Participants were instructed to pay attention to the episode and speculate the original storyline. While watching, they were encouraged to guess what the original temporal sequence of the story is, what causal links exist between events, and what characters are doing, thinking, and feeling. Each fMRI session comprised ten runs, each including the following three parts (**Fig. 1a**). Further details on fMRI data acquisition and preprocessing are also provided in Song et al^48^.

### Part I – Movie watching and aha button press

Participants were instructed to press an “aha” button whenever they understood something new or experienced a subjective feeling of insight while watching the movie. They were informed that there were no right or wrong answers and encouraged to respond without hesitation, even to the slightest feeling of realization. Specifically, we instructed participants to press the aha button when they:

- experience a subjective feeling of “Aha!”, or a sudden insight;
- thought they had grasped what was going on in the story;
- made sense of the past scene that they previously had not understood;
- had a better understanding of or had a speculation on the general plots, the temporal sequence of the events, and the causal relationship between events;
- had a better understanding of or had a speculation on characters’ behavior, thoughts, feelings, intentions, and relationships with other characters;
- resolved initial curiosity about any parts of the narrative, even partially;
- realized that their prior aha moments turned out to be incorrect (i.e., a feeling of oops).

### Part Ⅱ – Verbal explanation of aha moments

Following movie viewing, participants were presented with screenshots corresponding to their “aha” button presses, and were instructed to explain the reason for each press:

- What were your ‘aha!’ moments in these scenes?
- What have you newly understood/learned about the events and characters from watching these scenes?
- How did your thoughts about the events and characters change from watching these scenes? Participants were instructed to respond in detail, speak clearly and accurately (no mumbling), and minimize head and body movement. They were reminded that their voice was being recorded and to press a button to indicate completion. If multiple aha presses were made within a scene, the corresponding screenshots were presented together and numbered sequentially. Scenes with no aha presses were skipped.

### Part Ⅲ – Verbal report of character impressions

Participants finally reported their impressions of each of the four main characters in randomized order. They were instructed to report their thoughts about the characters, specifically:

- Describe the character’s personality traits
- Describe the character’s behavior, thoughts, feelings, intentions, motivations, or relationship with other characters
- What is your personal impression about the character?

Participants were reminded to speak aloud and clearly, provide as much detail as possible, and minimize head and body movement.

Following the fMRI session, participants completed a post-scan behavioral task, where they verbally recounted the narrative in its original order and rated the four main characters on a series of trait and impression scales. These data were collected but not analyzed in the present study.

### Coding of aha moment responses

Participants’ verbal explanations of aha moments were transcribed verbatim from in-scanner recordings. After reviewing these responses, the author H.S. defined eight mutually non-exclusive categories reflecting the types of insights: character, character relationship, memory retrieval, inference, temporal order, current event understanding, causal understanding, and “oops”. Authors J.K., R.M., and H.S. independently reviewed the transcriptions and labeled each aha moment into at least one of the eight categories (**Suppl. Table. 1**). Each “aha” button press could be categorized into one or more categories. In this paper, we focused on the “character” category (“character insight”), which is about comprehension of the character’s traits, behavior, thoughts or feelings. An aha moment was assigned to the “character” category if at least two annotators agreed on its classification.

### Nested mixed-effect model linking impression similarity with temporal distance

To examine how social impressions evolve over time within individuals, we built a nested mixed effects model relating impression similarity between every pair of runs with their temporal distance, with intercepts of four main characters, three scrambled-order groups, and subjects nested within groups included as random effects. Semantic meanings of verbal reports on character impressions were captured using Google’s Universal Sentence Encoder (USE), yielding 512-dimensional embedding vectors^108^. The use of this large language model in capturing the semantics of narratives has been validated in earlier studies^56–58^. Impression similarity was quantified as cosine similarity between USE-embeddings. Temporal distance reflects the number of runs separating two observations (e.g., the temporal distance between run 2 and run 5 is 3). A negative *β* coefficient for temporal distance indicates that impressions are less similar as their distance increases, reflecting gradual impression updating. We implemented the nested mixed effects model in R using the lmer function from the lme4 package.

### Intersubject representational similarity analysis (IS-RSA)

To test how the brain encodes social impressions, we conducted an IS-RSA analysis linking neural synchrony during movie-watching with similarity in reported impressions before and after the current movie-watching run (**Fig. 2a**). *Before*-movie impressions reflected participants’ prior thoughts and feelings about each character, allowing us to ask whether those who share similar initial impressions exhibited greater neural synchrony when later watching them onscreen. *After*-movie impressions represented impressions reported right after the movie blocks, testing whether individuals with higher neural synchrony during social perception subsequently developed more similar impressions. Impression similarity was assessed as cosine similarity of USE-embeddings from impressions on the same character, reported from different participant pairs within the same scrambled-order group. To account for the hemodynamic lag, neural time courses were shifted by 4 TRs (4.8s). For each ROI, neural synchrony was computed as the Pearson correlation between the voxel-average time series of two participants. Data from 7 participants were excluded from analysis due to excessive head motion (*N* = 3; > 10% total frames censored during movie-watching) or missing verbal transcripts (*N* = 4).

For each ROI, we calculated the Spearman correlation between between-participant impression similarity and neural synchrony across scenes. To ensure that these effects were not driven by between-group differences, we conducted the analysis separately within each of the three groups and computed the Fisher-*z* transformed mean *r*-value across the three groups. The resulting distribution of mean *r*-value was compared against zero with bootstrapping (10,000 iterations). A two-tail significance testing was performed, where *p* = (1+number of null abs(*r*-values) ≥ empirical abs(r)) / (1+number of permutations), and the resulting significance (i.e., *p*-values) was corrected for multiple comparisons across the 116 ROIs.

To identify regions in which neural synchrony tracked both before- and after-movie impressions, we combined the two IS-RSA analyses using Fisher’s method. For each ROI, *p*-values from the “before” and “after” analyses were first restricted to regions with consistent effect directions, and then combined as χ² = −2(ln p_before + ln p_after), which follows a chi-squared distribution with 4 degrees of freedom under the null hypothesis. We corrected for multiple comparisons across all 116 ROIs assessed in the IS-RSA analysis (corrected-*p* < 0.05).

### Neural synchrony mediates impression updating

To test whether shared neural responses during movie watching mediate the updating of social impressions across participants, we performed a causal mediation analysis, where we used pre-movie impression similarity as the predictor (T), neural synchrony as the mediator (M) and post-movie impression similarity as the outcome variable (Y).

For each ROI, we fit mediation models using the *mediation* package (v4.5) in R. The mediator model predicted ISC from pre-movie impression similarity while controlling for scene, subject pair, and group. The outcome model predicted post-movie impression similarity from both pre-movie impression similarity and neural ISC, again with scene, subject pair, and group included as categorical fixed effects. Mediation effects were estimated with nonparametric bootstrapping (1,000 iterations for the full 116-ROI analysis). We report the average causal mediation effect (ACME), average direct effect (ADE), total effect, and proportion mediated. ACME *p*-values were corrected for multiple comparisons across ROIs using the Benjamini–Hochberg false discovery rate (α = 0.05).

### Measuring neural pattern shifts near moments of character insight

To characterize the neural dynamics underlying insight moments, we examined whether neural representational patterns reconfigured around participants’ aha button presses—moments when participants experienced a sudden, new realization during movie watching. For each ROI, we quantified moment-to-moment neural pattern shifts as 1 - Pearson’s *r* between the voxel patterns in two consecutive TRs, such that higher values reflected greater representational change. These shifts were computed within a window from 5 TRs before to 3 TRs after each aha button press, following the selection of Song et al.^48^, with an expectation that participants would experience insight prior to pressing the aha button. For scenes where participants reported multiple aha moments, we averaged time courses of neural pattern shifts across the aha instances.

To test whether such neural pattern shifts were significantly above chance, we compared their magnitude to a null distribution generated by repeating the calculation during non-aha moments in the same movie-watching run (i.e., outside of the -5 to 3 TR window relative to any aha button press). For each ROI, we assessed the significance of pattern shifts at each TR within this window and performed correction for multiple comparisons both across the 9 TRs and the 116 examined ROIs. The number of significant ROIs was plotted in **Fig. 4d**. We performed a two-tail significance testing and conducted corrections for multiple comparisons across 116 ROIs and 9 assessed TRs.

### Linking neural pattern shifts with impression updating

Our central hypothesis is that impression updating is shaped by moments of insight into others’ traits, intentions, or mental states. To test this, we examined whether the magnitude of observed neural pattern shifts at character insight moments were associated with the degree of impression updating, quantified as 1 - cosine similarity of USE-embeddings of impressions from two consecutive runs (**Fig. 4b**). For each participant, we computed scene-wise impression updating, yielding an *n*_subject_× *n*_scene_ matrix. Similarly, for each ROI, we derived neural pattern shifts across participants and scenes, resulting in a corresponding *n*_subject_× *n*_scene_ matrix. Scenes where the corresponding subjects did not press the aha buttons were treated as NaNs, which was on average 4.5 out of 40 scenes (ranging from 4 to 6 scenes). There were 40 scenes (out of 48 scenes) because we excluded run 1 (5 scenes with no prior impressions) and run 7 (3 scenes with no aha categorization). To assess whether individuals with higher neural pattern shifts exhibit greater impression updating, for each scene, we computed Spearman correlations between the subject-wise vectors of neural pattern shifts and impression updating, and averaged the Fisher-*z* transformed correlations across scenes. We assumed a conservative significance testing, where the actual *r*-values were compared to a null distribution generated by repeating this analysis on non-insight moments in the same movie-watching run (1,000 iterations). We performed a two-tail significance testing, and corrected for multiple comparisons across 116 ROIs and 9 assessed TRs. We plotted ROIs that exhibited both significant neural pattern shifts and their correlation with subsequent impression updating in **Fig. 4e** for better interpretability of the results.

### Semantic characterization of aha and impression language

To assess whether character (consensus from 2 out of 3 raters; 45.8%) and non-character insights (none of 3 raters rated as character insight; 33.9%) are semantically different, we visualized the USE-embeddings of the aha transcripts in the UMAP 2-d space, and trained a logistic regression classifier with 5-fold stratified cross-validation (AUC; scaling within folds).

To test on what dimensions they were different, we tokenized each transcripts into lowercase words and calculated the proportion of tokens matching nine theory-driven, predefined semantic dictionaries (**Supplementary Table 4**): dispositional traits, mental states, social relationships, causal languages, individual-focus pronouns (e.g., he, she), group-focus pronouns (e.g., they), event/action terms^63–69^. We compared character with non-character insights across these categories using Wilcoxon rank-sum tests and FDR-corrected for multiple comparisons.

We derived two measures by averaging constituent categories. The *person model*, indexing representations of individual attributes, comprised dispositional traits, mental states, and individual focus. The *social-event model*, indexing external social or event structure, comprised social relationships, group focus, causal language, and event/actions. Model proportions were normalized to their sum, yielding a continuous partition between the two representations. We applied the same pipeline to social impression reports, yielding person- and social-event-model proportions across participants, characters, and runs.

### Predicting person vs social-event impression updating with aha counts

To examine whether runs with more insight moments tend to have more subsequent impression updating, we fitted linear mixed-effects models to predict the percentage change in the person vs social-event models in the impression reports from the number of aha button presses. We built one model for each of the four predictor–outcome combinations: person model ∼ character aha, situation model ∼ character, person model ∼ non-character, situation model ∼ non-character aha. Both predictor (aha count) and outcome (model proportion) were z-scored, such that the fixed-effect coefficient (*β*) reflects a standardized effect size. In these models, runs nested within subject and group were included as random variables: lme4: *y*₍z₎ ∼ *x*₍z₎ + (1 | group/subject/run)).

### Linking neural pattern shifts with person vs social-event updating in impressions

The analytical pipeline was identical to that used to link neural pattern shifts with semantic changes in impressions, except that the outcome variable was defined as the absolute change in model proportions rather than 1 − cosine similarity between embeddings. For each participant, r-values were first computed separately for each significant ROI and significant time point within the −5 to +3 TR window. These r-values were then averaged across all significant ROIs and time points to obtain a single summary measure for each condition. We then performed paired t-tests to compare person versus social-event model r-values within character or non-character insight ROIs, and independent t-tests to compare person or social-event model r-values between character and non-character insight ROIs.

## Data availability

The raw and preprocessed fMRI data of the socialaha dataset is publicly available on OpenNeuro https://openneuro.org/datasets/ds005658. The behavioral data are available at https://github.com/jinke828/socialaha/tree/main/data/beh. The intermediate brain data are available for convenience in replicating the findings: https://github.com/jinke828/socialaha/tree/main/data/brain.

## Code availability

Code to fully replicate the main analysis, plot the figures in the manuscript, and a step-by-step instruction to run these codes are available at: https://github.com/jinke828/socialaha/.

## Acknowledgments

We thank the Northwestern CTI, especially Rachael Young and Megan Dorn, for assisting with data collection. We thank Elizabeth Goldfarb and Nick Turk-Browne for their helpful comments on the project. Our work was supported by high-performance computing resources provided by Yale Center for Research Computing.

## Author contributions

Conceptualization: JK, MMC, MDR, YCL, HS

Methodology: JK, MMC, MDR, YCL, HS

Data collection: JK, RM, MDR, HS

Data curation, Formal analysis, & Visualization: JK, HS

Supervision: MMC, MDR, YCL, HS

Funding acquisition: MMC, MDR, YCL, HS

Writing—original draft: JK, MMC

Writing—review & editing: All authors

All the authors approved the final manuscript for submission.

