## Supplementary information for "Neural dynamics of updating social impressions during movie watching"

Jin Ke *et al*

**This PDF file includes:**

Figs. S1 to S9

Tables. S1 to S4

**Supplementary Figures**

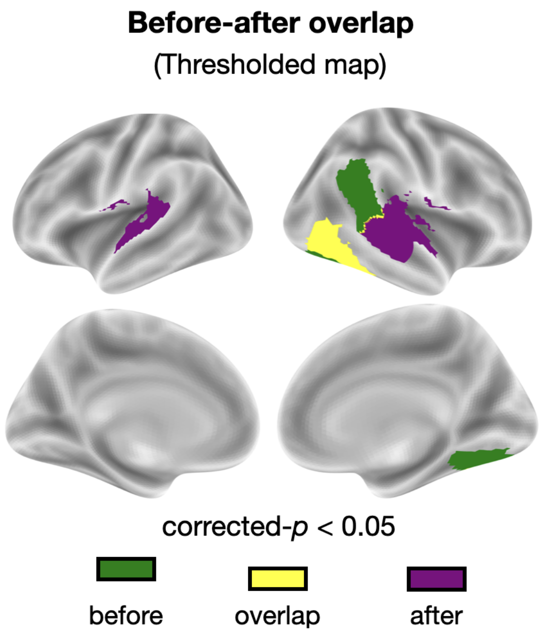


**Supplementary Figure 1.** Regions where synchrony increases with greater similarity in before-movie impressions (green; the right temporoparietal junction, angular gyrus, superior temporal sulcus and inferior temporal cortex) or after-movie impressions (purple; bilateral medial temporal gyrus, the right superior temporal sulcus, superior temporal gyrus, auditory and inferior temporal cortex. The yellow area indicates overlapping regions between before and after (inferior temporal cortex). Only regions surviving correction for multiple comparisons across all ROIs (corrected-$p$ $< 0.05$, two-tailed test) are shown.

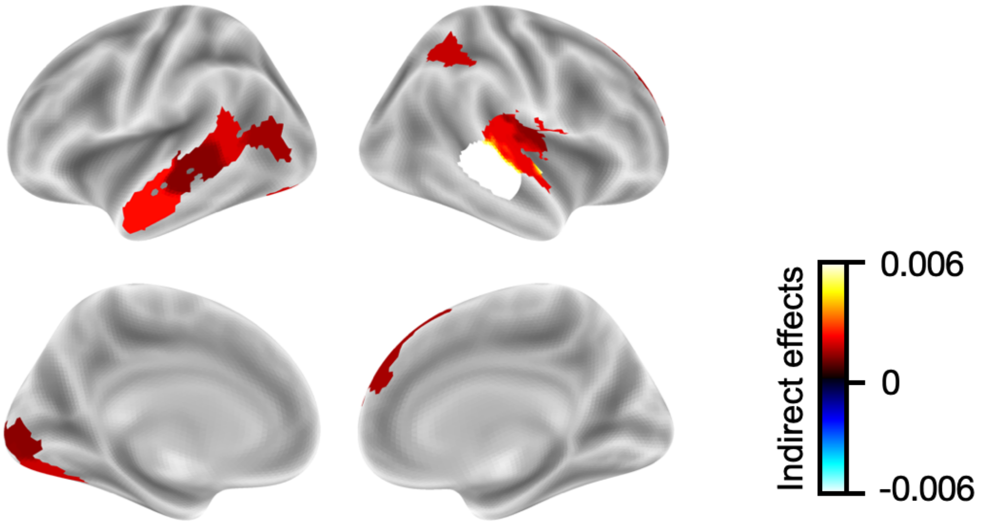


**Supplementary Figure 2.** Regions whose neural synchrony during movie-watching exhibit significant (uncorrected-$p < 0.05$) or marginal (uncorrected-$p < 0.10$) mediation effect on the relationships between before- and after-movie impressions, before correcting for multiple comparisons.

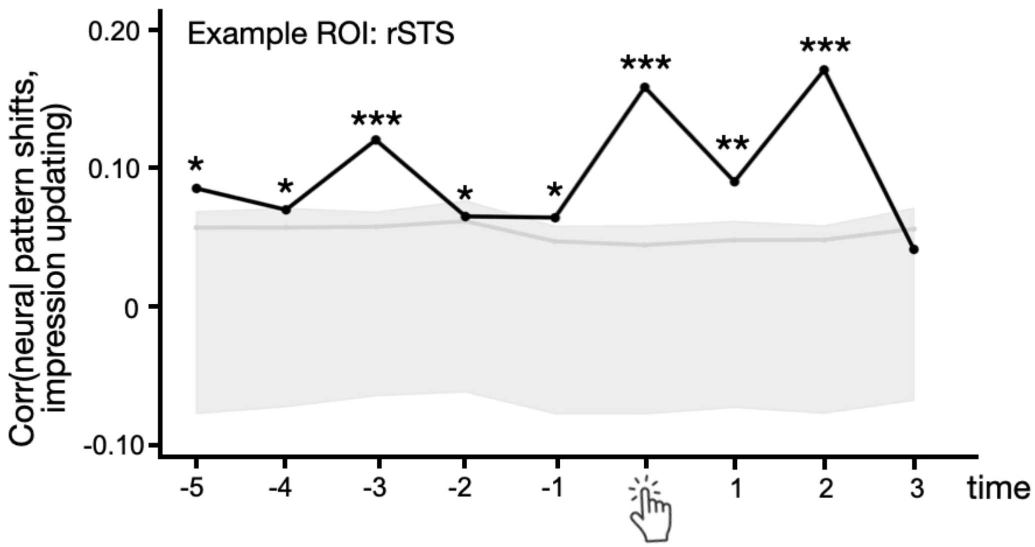


**Supplementary Figure 3.** Correlations between neural pattern shifts in the right superior temporal sulcus and impression updating as a function of time relative to aha button presses. The gray area represents a null distribution of these correlations from non-insight moments. The gray line indicates the top 5% *r*-value in the null distribution. The asterisks indicate statistical significance comparing the *r*-values to the null distribution with one-tailed tests at each TR (before correction). *: $p < 0.05$, **: $p < 0.01$, ***: $p < 0.001$.

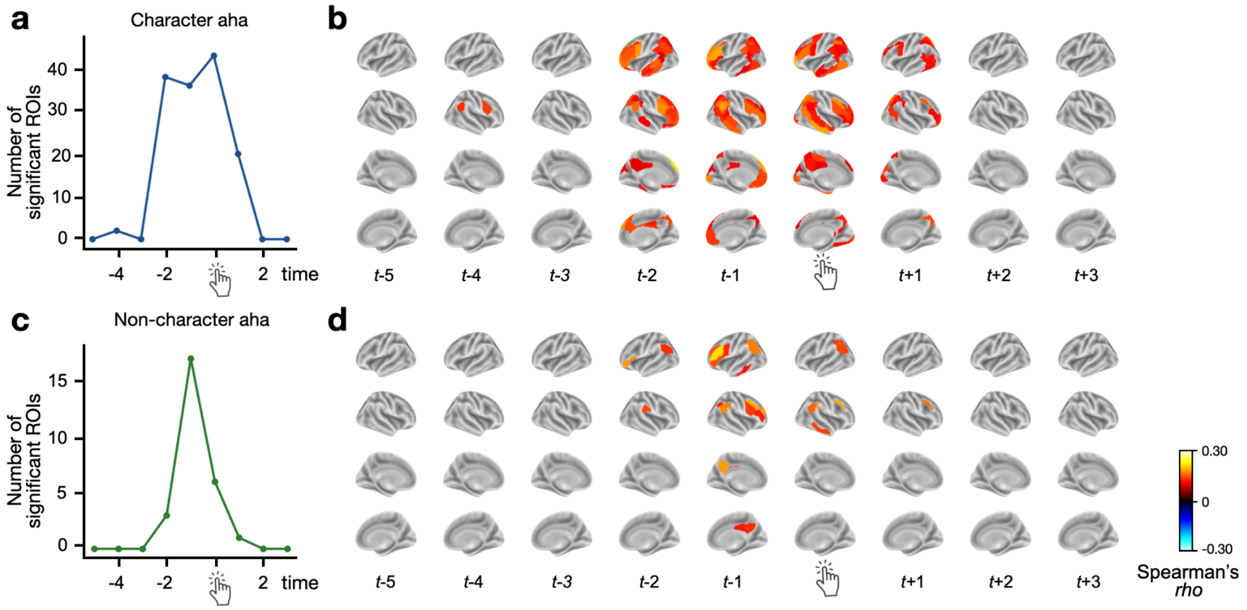


**Supplementary Figure 4.** Neural pattern shifts at insight moments. **(a)** Number of ROIs with significant neural pattern shifts near *character aha* moments as a function of time. **(b)** Whole-brain neural pattern shifts as a function of time relative to *character aha* button presses. Brain maps plot the *z*-statistics comparing neural pattern shifts near character aha moments with those during non-aha moments. Regions with significant neural pattern shifts were plotted, corrected for multiple comparisons across 116 ROIs and 9 TRs (corrected-$p < 0.05$) This figure is identical to Fig. 4d. **(c)** Number of ROIs with significant neural pattern shifts near *non-character aha* moments as a function of time. **(d)** Whole-brain neural pattern shifts as a function of time relative to *non-character aha* button presses. Brain maps plot the *z*-statistics comparing neural pattern shifts near non-character aha moments with those during non-aha moments. Regions with significant neural pattern shifts were plotted, corrected for multiple comparisons across 116 ROIs and 9 TRs (corrected-$p < 0.05$)

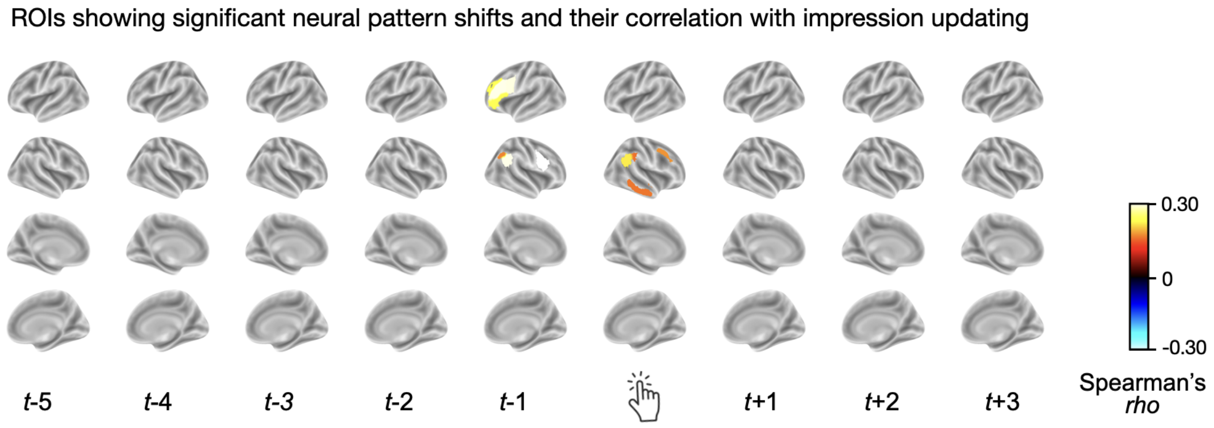


**Supplementary Figure 5.** Neural pattern shifts and correlation with impression updating during *non-character aha* moments. ROIs showing both significant neural pattern shifts and their correlation with impression updating. Significance was corrected for multiple comparisons across 116 ROIs and 9 TRs.

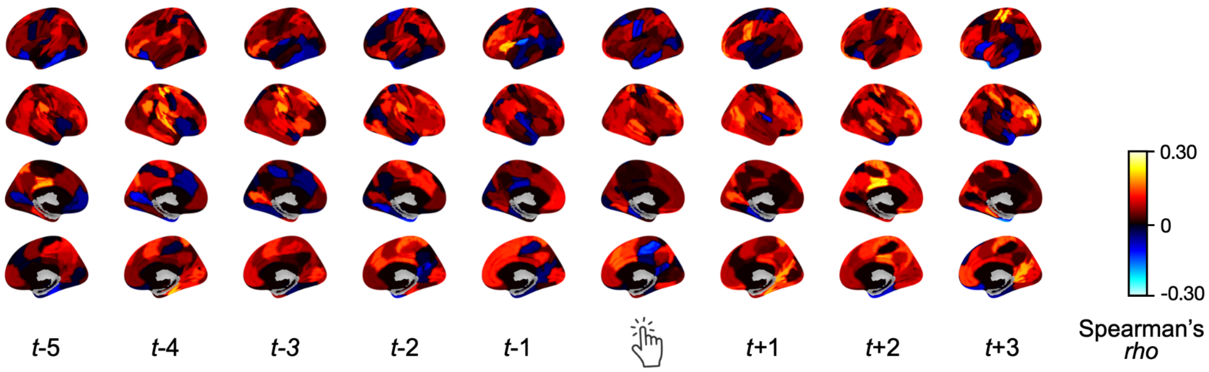


**Supplementary Figure 6.** Raw brain maps of correlations between neural pattern shifts near character insight with impression updating.

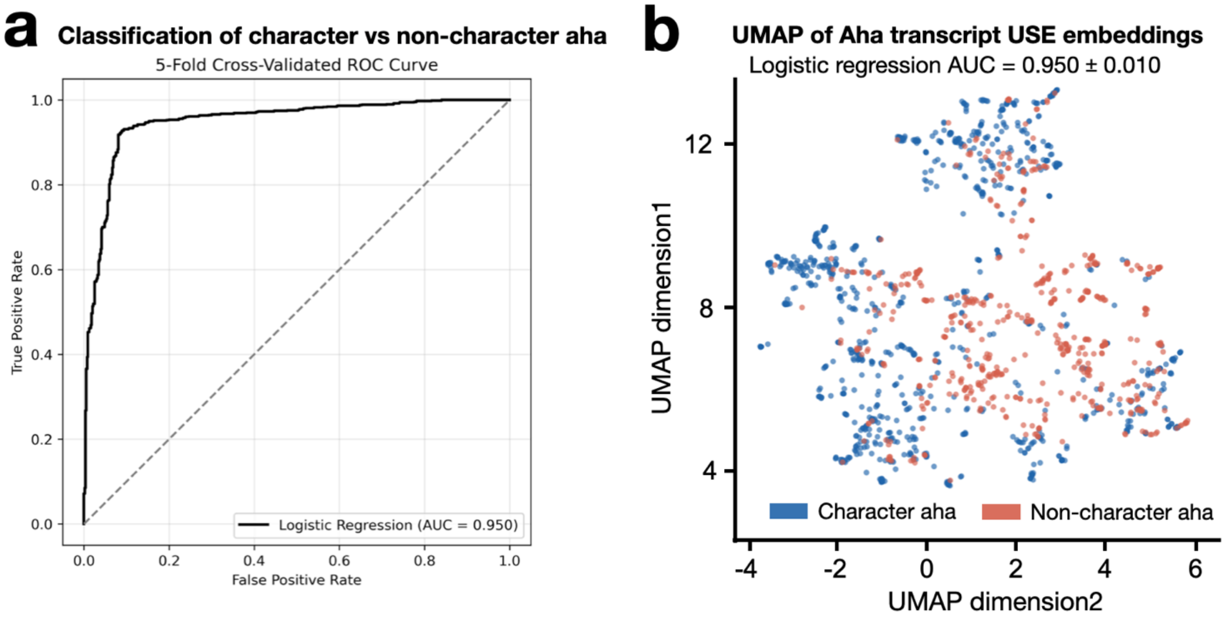


**Supplementary Figure 7. (a)** Cross-validated classification of USE embeddings of character vs. non-character aha transcripts. Receiver operating characteristic (ROC) curve for the logistic regression classifier evaluated using 5-fold cross-validation. The solid black line represents the average ROC curve across folds, showing the trade-off between true positive rate (sensitivity) and false positive rate (1 − specificity). The dashed diagonal line indicates chance-level performance. **(b)** Universal Sentence Encoder (USE)-embeddings of aha transcripts projected onto the first two dimensions of the Uniform Manifold Approximation and Project (UMAP) space. Each dot represents an aha verbal report from a single participant, run, and character.

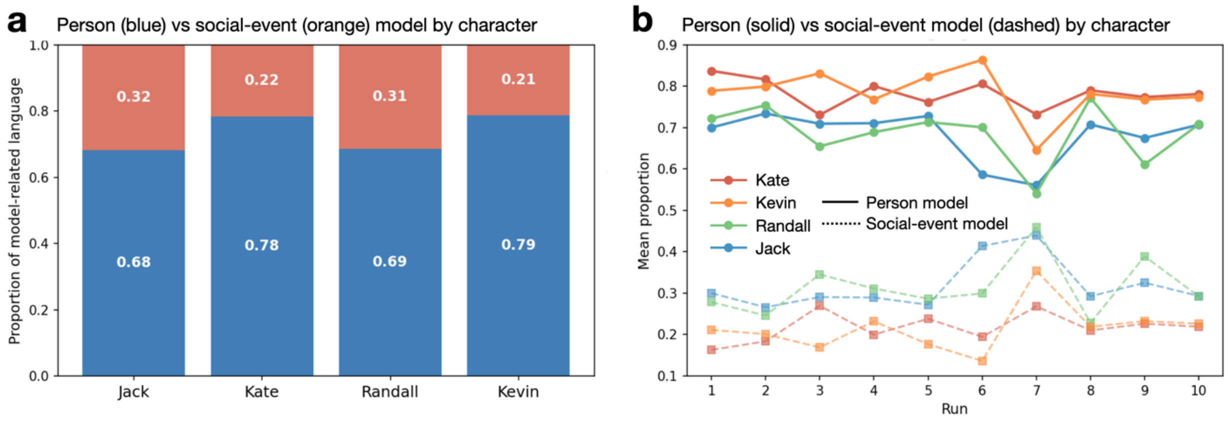


**Supplementary Figure 8.** Language used for reporting character impressions consistently showed higher person model proportions than social-event model proportions across characters and runs. **(a)** Proportions of person vs social-event model language for each character. Blue indicates person model, orange indicates social-event model. **(b)** Proportions of person vs social-event model language for each character as a function of run. Solid line indicates person model, dashed line indicates social-event model.

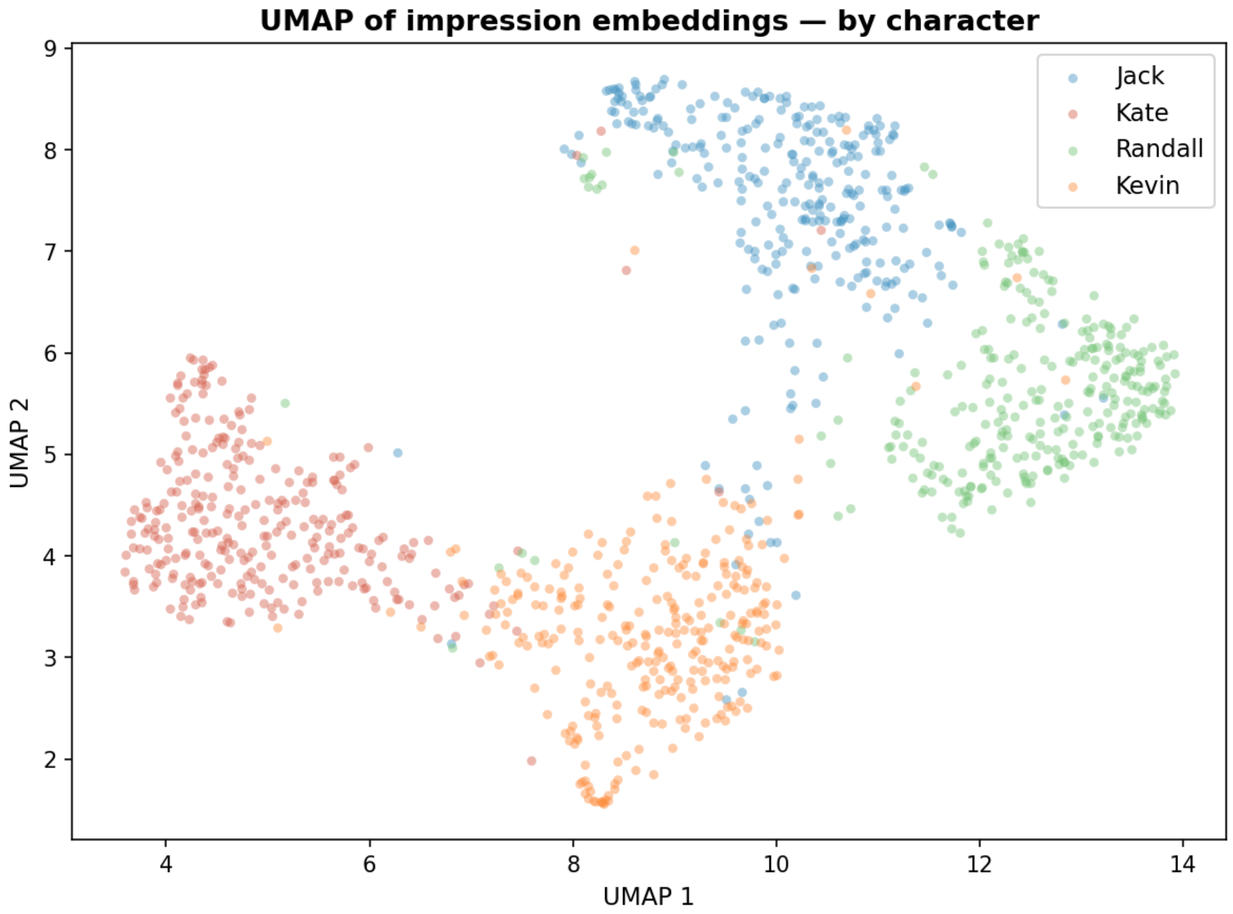


**Supplementary Figure 9.** Universal sentence encoder (USE) embeddings of impression transcripts projected onto the first two dimensions of the Uniform Manifold Approximation and Project (UMAP) space. Each dot represents a verbal impression report from a single participant, run, and character.

### Supplementary Tables

**Table. S1** Definitions of each aha category and their proportions among all aha moments. Verbal explanations for each of the aha button presses from 32 participants were coded into one or more categories (i.e., categories were not mutually exclusive). Categories were derived from participants' responses and applied independently by three raters. For each participant, we calculated the percentage of aha moments assigned to each category (with agreement from >= 2 raters) and then averaged these percentages across participants. In the present study, analyses focused on character-related aha moments. To ensure data quality, we included only those instances that were labeled as "character" by at least two of the three raters (i.e., majority consensus). The non-character aha was defined as the aha moments where none of the three raters labeled as character aha.

| **Aha category** | **Description** | **Mean proportions (SD)** at least 2 raters |
|---|---|---|
| Memory retrieval | Explicit or implicit mention of past events | 47.2 (19.2) % |
| Character | Comprehension of the character's traits, behavior, thoughts or feelings | 45.8 (19.2) % |
| Current scene understanding | Comprehension or descriptions of what is happening in the current event | 41.2 (17.8) % |
| Character relationship | Comprehension of the relationship between characters | 19.1 (7.6) % |
| Temporal order | Comprehension of the original temporal order of the events | 14.5 (12.6) % |
| Inference | Inference on narrative contents or connection links between events; Prediction of future events | 8.2 (6.4) % |
| Causal understanding | Comprehension of causal structure of the narrative, or causal relationship between events. | 7.3 (5.1) % |
| Oops | Realizing prior comprehension had been incorrect and/or correcting for it. | 4.8 (4.6) % |

**Table. S2** Segmentation of the full *This Is Us* episode into 48 events, with each event's brief description, featured main character(s), and duration. Segmentation was based on the director's cut, with each scene depicting a single character's narrative except for scenes labeled "Kevin & Kate". These 48 scenes were scrambled in three different orders and presented across 10 fMRI runs, with N=12 per scrambled-order group.

| Scene id | Description | Character | Length (s) | Scene id | Description | Character | Length (s) |
|---|---|---|---|---|---|---|---|
| 1 | Opening, Birthday towel | Jack | 71 | 25 | Leave office | Randall | 15 |
| 2 | Birthday cake in the fridge, Scale | Kate | 34 | 26 | Father's house (outside) | Randall | 87 |
| 3 | Work, Good news email, Happy birthday | Randall | 26 | 27 | Father's house (inside) 1 | Randall | 80 |
| 4 | Bedroom with two girls, I am 36 | Kevin | 31 | 28 | Father's house (inside) 2 | Randall | 43 |
| 5 | Big three, Terrible things to mommy | Jack | 30 | 29 | Behind the scene with Alan Thicke | Kevin | 33 |
| 6 | Click good news email | Randall | 9 | 30 | A scene with Alan Thicke | Kevin | 59 |
| 7 | Scale naked, Falls back | Kate | 18 | 31 | Alternative lighter version | Kevin | 54 |
| 8 | Kiss, My water just broke | Jack | 16 | 32 | Meltdown 1 | Kevin | 54 |
| 9 | The Challenger story | Kevin | 67 | 33 | Meltdown 2: Shame on you, I quit | Kevin | 67 |
| 10 | Bathroom floor, How did we get here? | Kevin & Kate | 50 | 34 | Rebecca in labor, Something's not right | Jack | 44 |
| 11 | Bathroom floor, I ate my dream life away | Kevin & Kate | 72 | 35 | Father meets family | Randall | 51 |
| 12 | Hospital room, six weeks early birth | Jack | 39 | 36 | Cracking up | Randall | 38 |
| 13 | Deep breath, High risk pregnancy | Jack | 54 | 37 | Date at a restaurant | Kate | 44 |
| 14 | Dr. Katowski, Best of the best, Bad joke | Jack | 66 | 38 | Invite in? | Kate | 58 |
| 15 | Man-ny set, Breastfeed | Kevin | 38 | 39 | Pretty picture | Kate | 47 |
| 16 | Ryan Gosling would never do this crap | Kevin | 54 | 40 | Toby meets Kevin, meltdown news | Kevin & Kate | 55 |
| 17 | Soccer field with wife and two daughters | Randall | 35 | 41 | Doctor shares the news | Jack | 86 |
| 18 | I found him | Randall | 48 | 42 | Doctor's story 1 | Jack | 81 |
| 19 | Throwing food away, Dog shit | Kate | 29 | 43 | Doctor's story 2 | Jack | 95 |
| 20 | Group therapy 1 | Kate | 33 | 44 | I'm dying. Foster parents? | Randall | 97 |
| 21 | Group therapy 2 | Kate | 58 | 45 | What am I going to do? | Kevin & Kate | 72 |
| 22 | Coffee, You want to be fat friends? | Kate | 58 | 46 | Jack and the fireman at the hospital | Jack | 61 |
| 23 | Breathe, I like you two | Jack | 40 | 47 | Everyone | Everyone | 71 |
| 24 | Conversation, Three healthy babies | Jack | 77 | 48 | The Big Three, Ending | Jack | 52 |

**Table. S3** Regions whose neural synchrony during movie-watching exhibit significant (uncorrected-$p < 0.05$) or marginal (uncorrected-$p < 0.10$) mediation effect on the relationships between pre- and post-movie impressions. Here we report the ROI label from the Schaefer 100 atlas, and the statistics from the causal mediation model, including the averaged causal mediation effect (ACME) and its 95% confidence interval, and the corresponding *p*-values (uncorrected).

| **ROI name** | **ACME** | **95% CI** | **uncorrected-*p*** |
|---|---|---|---|
| 17Networks_LH_VisCent_ExStr_1 | 0.0017 | [0.0001, 0.0041] | 0.032* |
| 17Networks_LH_VisCent_ExStr_3 | 0.0012 | [-0.0001, 0.0031] | 0.082† |
| 17Networks_LH_DorsAttnA_SPL_2 | 0.0013 | [-0.0001, 0.0033] | 0.092† |
| 17Networks_LH_DefaultB_Temp_1 | 0.0023 | [0.0006, 0.0047] | 0.004** |
| 17Networks_LH_DefaultB_Temp_2 | 0.0011 | [-0.0001, 0.0032] | 0.086† |
| 17Networks_LH_TempPar_1 | 0.0019 | [0.0003, 0.0041] | 0.008** |
| 17Networks_RH_SomMotB_S2_2 | 0.0012 | [0.0000, 0.0029] | 0.06† |
| 17Networks_RH_ContA_IPS_1 | 0.0016 | [0.0002, 0.0037] | 0.024* |
| 17Networks_RH_DefaultB_PFCd_1 | 0.0014 | [-0.0001, 0.0034] | 0.074† |
| 17Networks_RH_TempPar_2 | 0.0060 | [0.0030, 0.0099] | < 0.001*** |

**Table. S4** Word dictionaries used to quantify semantic content in aha and impression transcripts. Words were drawn from prior theory-driven lexicons in social cognition and language research, covering categories such as dispositional traits, mental states, social relationships, causal language, individual-focus pronouns, group-focus pronouns, and event/action terms. Transcripts were tokenized, and the proportion of words matching each category was computed. Categories were combined into two measures: the person model (traits, mental states, individual focus) and the social-event model (relationships, group focus, causal language, events).

| Category | Dictionary |
|---|---|
| Mental state (Tamir et al., 2016) | affection, agitation, alarm, alertness, amazement, ambivalence, amusement, anger, annoyance, anticipation, anxiety, apathy, appreciation, apprehension, attention, awareness, awe, belief, bewilderment, bias, bitterness, boredom, calmness, certainty, cheerfulness, cognition, concern, confusion, consciousness, contemplation, contempt, contentment, curiosity, decision, delight, depression, desire, despair, disappointment, disbelief, disgust, distress, distrust, dominance, dread, dreaminess, drowsiness, earnestness, ecstasy, elation, embarrassment, emotion, empathy, enjoyment, enthusiasm, envy, exasperation, excitement, exhaustion, expectation, fascination, fatigue, feeling, frenzy, friendliness, frustration, fury, gloominess, guilt, hallucination, happiness, horror, humiliation, humor, hunger, hypnosis, hysteria, imagination, impatience, indecisiveness, indifference, insanity, inspiration, intention, interest, intrigue, irritation, jealousy, judgment, laziness, lethargy, loneliness, lust, melancholy, memory, misery, mortification, nervousness, objectivity, opinion, optimism, outrage, pain, panic, patience, peacefulness, pensiveness, pity, planning, playfulness, pleasure, prejudice, preoccupation, pride, rage, reason, regret, relaxation, relief, remorse, resentment, sadness, satisfaction, self-consciousness, self-control, self-pity, serenity, seriousness, shame, shock, skepticism, sleepiness, sorrow, stress, stupor, subordination, surprise, suspicion, sympathy, terror, thirst, tiredness, torpor, trance, transcendence, uncertainty, uneasiness, unhappiness, vengeance, wakefulness, warmth, weariness, woe, worry |
| Dispositional traits (Thornton and Mitchell, 2018) | warmth, competence, agency, experience, trustworthiness, dominance, openness, conscientiousness, extraversion, agreeableness, neuroticism, attractiveness, intelligence, kind, generous, selfish, ambitious, insecure, confident, shy, honest, dishonest, caring, cold, warm, jealous, loyal, responsible, determined, stubborn, sensitive, emotional, successful, talented, smart, proud, humble, bitter, guilty, ashamed, strong, weak, depressed, anxious, upset, angry, frustrated, passionate, devoted, dedicated, driven, identity, personality, character, nature |
| Individual focus | he, him, his, she, her, he's, she's |
| Social relationship | father, dad, mother, mom, son, daughter, brother, sister, sibling, siblings, wife, husband, partner, friend, family, parent, parents, child, children, baby, babies, couple, triplet, triplets, adopted, biological, stepfather, stepmother, marriage, relationship, together, married |
| Group focus | they, them, their, they're, themselves |
| Event/Action | happened, occurs, occurred, found, discover, discovered, revealed, reveal, learned, learn, met, meet, told, tell, showed, show, died, dead, born, left, arrived, arrive, went, came, saw, seen, heard, did, done, got, gotten, took, taken, gave, given, brought, sent, lost, won, fell, broke, ran, turned, moved, started, ended, began, finished, decided, chose, chosen, tried, failed, succeeded |
| Causal language | because, therefore, thus, hence, explains, explanation, reason, caused, resulted, consequently, since, due, why, leads, led, means, implies, suggest, suggests, indicating, indicates |
